# Palaeontological evidence for community-level decrease in mesopelagic fish size during Pleistocene climate warming in the eastern Mediterranean

**DOI:** 10.1101/2022.10.04.510798

**Authors:** Konstantina Agiadi, Frédéric Quillévéré, Rafał Nawrot, Theo Sommeville, Marta Coll, Efterpi Koskeridou, Jan Fietzke, Martin Zuschin

## Abstract

Mesopelagic fishes are an important element of marine food webs, a huge, still mostly untapped food resource, and great contributors to the biological carbon pump, whose future under climate change scenarios are unknown. The shrinking of commercial fishes within decades has been an alarming observation, but its causes remain contended. Here, we investigate the effect of warming climate on mesopelagic fish size in the eastern Mediterranean Sea during a glacial–interglacial–glacial transition of the Middle Pleistocene (marine isotope stages 20–18; 814–712 Kyr B.P.), which included a 4 °C increase of global seawater temperature. Our results based on fossil otoliths show that the median size of lanternfishes, one of the most abundant groups of mesopelagic fishes in fossil and modern assemblages, declined by ~35% with climate warming at the community level. However, individual mesopelagic species showed different and often opposing trends in size across the studied time interval, suggesting that climate warming in the interglacial resulted in an ecological shift toward increased relative abundance of smaller-sized mesopelagic fishes due to geographic and/or bathymetric distribution range shifts, and the size-dependent effects of warming.

## Introduction

Climate change affects fish size, fitness, abundance, and distribution (1–3), with devastating anticipated socioeconomic impacts (4). Within the euphotic zone, where most human activities take place and seawater temperature is directly regulated by local climatic conditions, average fish size has been predicted to decrease by 14–24% by the year 2050 (5). Climate change is expected to have a significant impact on the mesopelagic zone as well (i.e., the part of the water column in the world’s oceans between 200 and 1000 m), by rapidly displacing isotherms (6): models predict a resulting expansion and shallowing of the deep scattering layers of the ocean water column, where most mesopelagic organisms live, leading to homogenization of their community composition and changes in mesopelagic biomass (7), and these predictions are so far confirmed by observations (8). Fishes are a dominant component of the mesopelagic communities with an estimated biomass of 2-19.5 Gtn, approximately 100 times greater than that of the total global annual fishery catches (9,10). Mesopelagic fishes are an important component of the ecosystem today and in the past (11–15), having a significant contribution to the biological carbon pump (16) through their light-controlled diel vertical migrations (17). They usually occupy the mesopelagic realm during daytime and migrate to surface waters at night to feed, functioning as a trophic link between primary consumers and megafauna (18,19). In fact, they help maintain ecosystem stability under environmental change (20), and their night-time movement upwards into surface waters in order to feed while avoiding predators leads to a downward flux of organic carbon from the productive euphotic zone into the deeper parts of the ocean (21–23). Due to this function, any adverse effects of climate warming on mesopelagic fishes directly impacts the oceans’ ability to sequester carbon from the atmosphere to the deeper parts of the ocean. Moreover, due to the current state of global food security, they are an important potential food resource (9,24,25). Indeed, mesopelagic fishes are a rich, targeted food resource, with 2.68 million tons of reported catches between 1950 to 2018 globally, even without proper exploitation efforts (24), and much higher estimated total biomass (26).

Body size is a key biological trait, which plays a significant role in controlling the structure and functioning of marine ecosystems (27), and it is strongly affected by ambient water temperature (28,29). Evaluating and predicting the long-term effects of climate change on size structure of modern fish assemblages is challenging, because they are difficult to disentangle from the impacts of other anthropogenic stressors like size-selective harvesting (30,31). Moreover, fishery and scientific survey data rarely encompass more than a few decades and are often biased towards commercially important species. The fossil record provides a rich archive of major environmental perturbations of the geological past and their paleoecological consequences, allowing us to track biotic responses to natural climatic shifts on timescales well beyond the limits of ecological monitoring (32). Here, we quantify the effect of the Pleistocene climatic variability on the size structure of the eastern Mediterranean mesopelagic fish assemblages across the time interval 814–712 Kyr B.P. corresponding to the marine isotope stages (MIS) 20–18: MIS 20 glacial (814–761 Kyr B.P.), MIS 19 interglacial (761–757 Kyr B.P.) and MIS 18 glacial (757–712 Kyr B.P.; 33,34). Our study is based on fossil otoliths from a unique hemipelagic sedimentary succession of this age exposed on the island of Rhodes in the eastern Mediterranean.

In contrast to present-day ecosystems, Pleistocene marine ecosystems were affected by the severe climatic oscillations of the glacial and interglacial periods, which can be used as analogues of current and forecasted climate warming situations without the confounding effects of the multiple anthropogenic stressors impacting present-day ecosystems. The Early–Middle Pleistocene Transition encompasses our target time interval MIS 20–18 and was characterized by important changes in Earth’s climate, when the duration of climate oscillations increased, leading to the growth of the Northern Hemisphere ice-sheets during glacial periods and a shift to the modern climatic regime with stronger climatic fluctuations (33,35). The interval MIS 20–18 involved a global sea surface temperature increase of about 4 °C (36,37), also expressed in the Mediterranean (38,39), which took place over a few thousands of years (MIS 20–19 deglaciation 786–789 Kyr B.P.; 40–41) and is in line with the IPCC-predicted mean surface temperature increase under the high greenhouse gas emissions scenario that is expected to disrupt the marine food-web structure irreversibly (1, 42–44). In the eastern Mediterranean, previous studies have shown that the mesopelagic fish fauna was indeed affected by the Pleistocene climatic perturbations: North Atlantic and Arctic fish species, whose geographic distribution included the eastern Mediterranean during glacial periods after 1.5 Ma B.P., became extirpated by the subsequent interglacials (11). Although the timescale over which the MIS 20–19 Pleistocene deglaciation took place was longer than the modern unprecedented climate change by one order of magnitude (40,41), the MIS 20–18 interval allows evaluating the potential long-term impact of a climate change on marine ecosystems and thus constrain the range of the possible future biotic responses on the on-going warming.

We reconstructed changes in body size and composition of Pleistocene mesopelagic fish assemblages using fossil otoliths. Fish otoliths are aragonitic incremental biomineralisates with species-specific morphology (43) that are commonly preserved as fossils in marine sediments, and whose assemblages faithfully record past fish faunas (14,46,47). Otolith size correlates with fish size through species-specific functions (48), although, in rare cases, otolith growth may become decoupled from fish growth (49). Here, we used otolith length–fish length or otolith width–fish length, and fish length–weight functions derived from present-day fishes to estimate Pleistocene fish sizes during MIS 20, 19 and 18. Changes in average size at the scale of entire assemblages can result from processes occurring at different levels of biological organization: from individual to community scales (28,29). Therefore, we also tracked changes in body size and relative abundances of individual species to evaluate the relative importance of size shifts within population and of changes in community composition in driving the observed body size patterns.

## Material and Methods

The samples were obtained on the island of Rhodes in the south-eastern Aegean Sea, in the eastern Mediterranean (Figure 1). Rhodes is part of the Hellenic forearc and has experienced intense vertical tectonic movements during the past 2 Myrs, leading to the deposition at bathyal depths and recent uplift of Early and Middle Pleistocene hemipelagic sediments, now exposed onshore along its eastern coast (50–53). The occurrence of such Pleistocene deep-water sediments accessible on land is unique for the eastern Mediterranean, providing a reference point for studying the Early and Middle Pleistocene climates in this region (53): there are no other deep-sea sediments covering this time interval cropping out across the region (51,54–56). We sampled the marl levels of the Lindos Bay Formation in the lower part of Lardos section on Rhodes (N 37°17’48’’, E 27°8’4’’), LR15–19, LR19–27, and LR>27, corresponding to MIS 20, MIS 19, and MIS 18, respectively (57).

**Figure 1.**
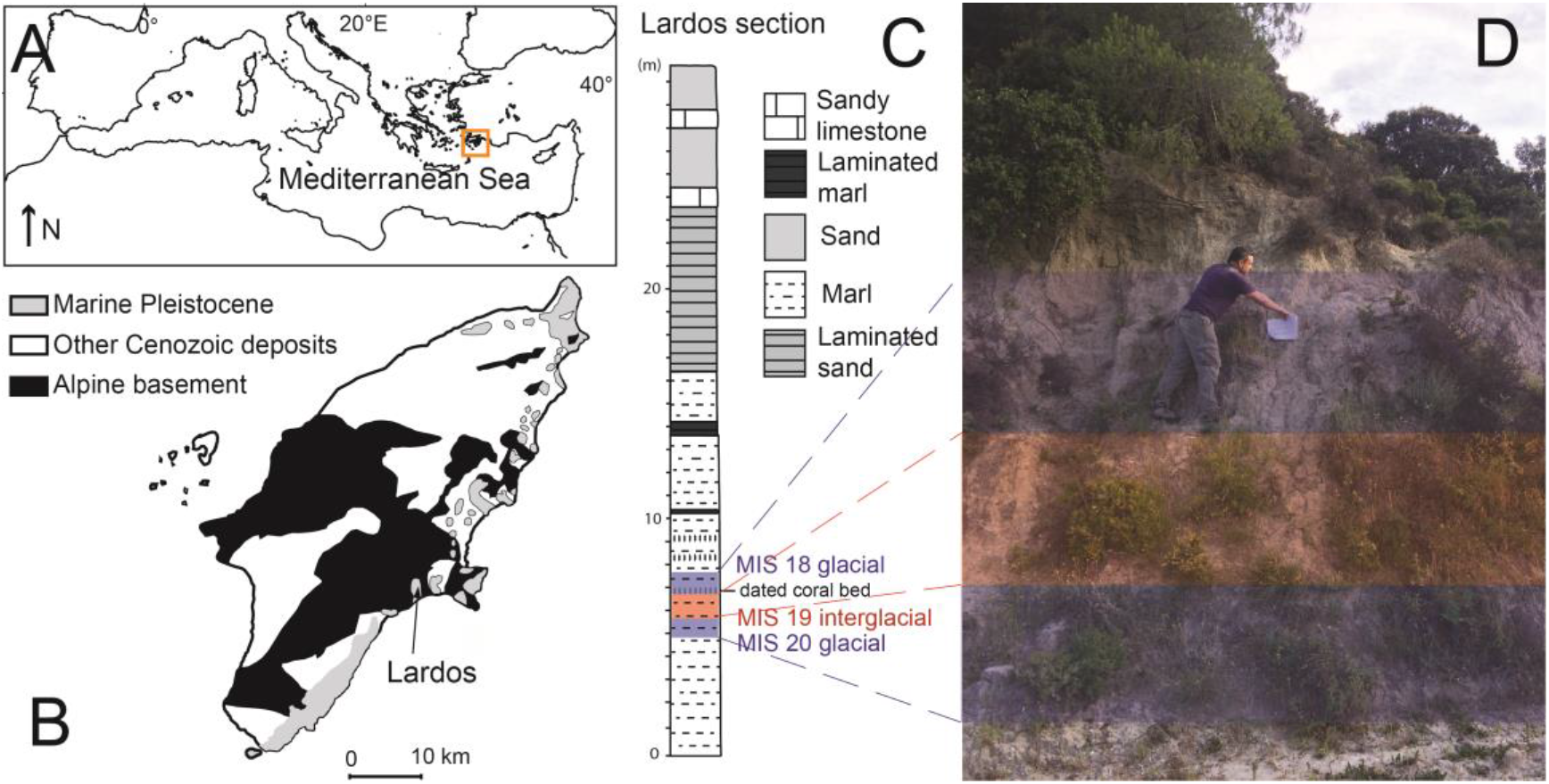
Sampling location: A. Map of the Mediterranean Sea; B. simplified geologic map of Rhodes, showing the location of the studied Lardos section (modified after (53)); C. lithological column of the Lardos section, indicating the studied MIS20–18 interval; and D. photograph of the sampled part of the outcrop.

The sediments of the Lardos section were deposited continuously with an average sedimentation rate of 1.9 cm/Kyr (57) and show uniform lithologies. The palaeodepth in the study area exceeded 200 m for both MIS 20 and MIS 19, but became slightly shallower during MIS 18, as indicated by the greater abundance of goby otoliths (11). The sediment samples were diluted in water and sieved with a 250-μm mesh sieve, and then dried in an oven. The sagittal otoliths (henceforth referred to simply as ‘otoliths’) were handpicked from the residues and identified based on the morphological characteristics (45).

Each otolith was photographed, and its length and width were measured (Figure S1) using the microscope ZEISS SteREO Discovery V20 and the software ZEN (ZEISS Efficient Navigation). Since most of the identified species are extant, we estimated their weight using modern empirical otolith length/width–fish length (Table S1) and the fish length–weight functions developed based on fish shape (58,59). The climatic-zone affinity and the habitat (pelagic or demersal) of the identified species were obtained from AquaMaps (60) and Fishbase (61), respectively (Table S2). We distinguished: (1) warm-water species, occupying (sub)tropical latitudes and (2) cold-water species, occurring in temperate and/or subpolar latitudes.

We examined changes in the frequency distributions of the otolith length, and fish lengths and weights across the three time intervals for the entire assemblages. To depict the main drivers of these patterns, we traced shifts in median fish weight and relative abundances within the most abundance species, individual families and climatic-affinity groups. As the size-frequency distributions were strongly right-skewed, all analyses were performed on log-transformed data. We used a non-parametric Kruskal-Wallis test, followed by a pairwise Wilcoxon test with a-posteriori Bonferroni correction to compare median weight between the three assemblages and estimate 95% confidence intervals around this parameter using a bootstrap procedure with 10,000 iterations. To test for differences in otolith preservation that could affect the interpretation of our results, we quantified and statistically compared the otolith preservation state following a previously presented approach (62). Specimens that could not be identified at least to family level were included in the otolith preservation analysis, but excluded from size calculations. All analyses were performed in R (version 4.1.2) (R Development Core Team 2021).

## Results

We identified and estimated the fish length and weight from the length or width of 1960 otoliths from the three time intervals: 655 otoliths (97.05% of the assemblage) from MIS 20, 1022 otoliths (98.46%) from MIS 19, and 283 otoliths (93.40%) from MIS 18 (Table S3). The otolith length ranged from 0.38 to 8.69 mm (median of 1.40 mm), and the otolith width ranged from 0.41 to 4.68 mm (median of 1.34 mm).

Overall, the fish in the three assemblages are small, with a median length of 3.22 cm (ranging from 0.82 to 10.07 cm) and a median weight of 0.27 g (ranging from 0.002 to 106.13 g) (Figures 2 and S3; Table S3). The small size of the fish is expected, given the nature of the otolith fossil record and the sampling method (63). The transition from MIS 20 glacial to MIS 19 interglacial was associated with 36% decrease in the assemblage-level median fish weight (from 0.36 g to 0.23 g), followed by 73% increase in the subsequent MIS 18 glacial period (to 0.40 g; Kruskal-Wallis test *χ*^2^ = 65.104, df = 2, *p* < 0.001; pairwise Wilcoxon test: *p* < 0.001 for both MIS 20–MIS 19 and MIS 19–MIS 18; Figure 2; Table S3).

**Figure 2.**
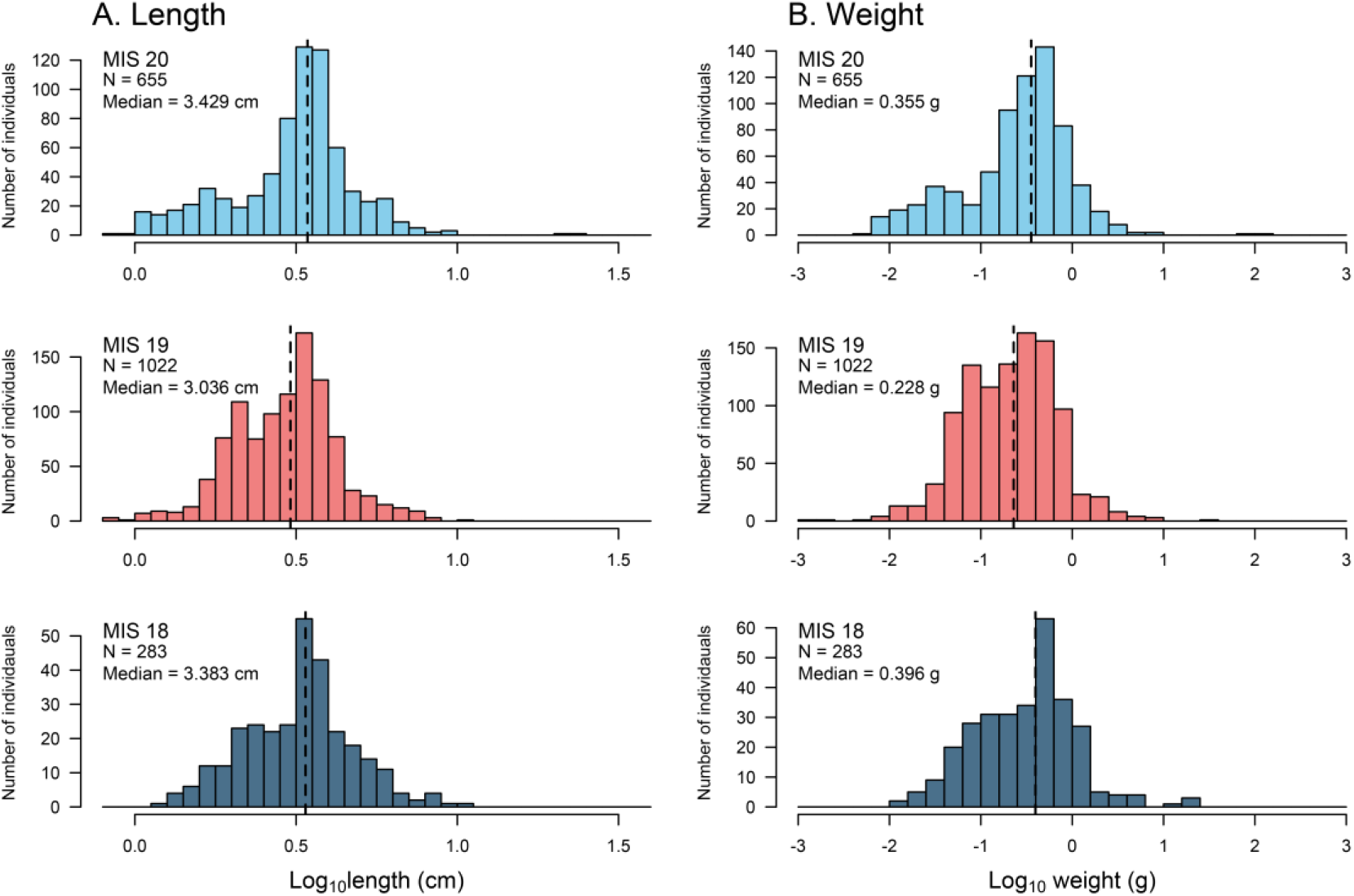
Fish length (A) and weight (B) frequency distributions (log-transformed) during MIS 20 (glacial), MIS 19 (interglacial), and MIS 18 (glacial). The median fish size at the assemblage level is smaller during the MIS19 interglacial.

On the genus and species level, lanternfishes’ sizes show contrasting trends (Figure 3; Table S4). The subtropical *Lobianchia dofleini*, the only representative of its genus, has similar median size throughout the studied interval with only a slight increase during MIS 19, as does the temperate *Hygophum benoiti*. However, the abundance of *L. dofleini* increases relative to *H. benoiti* in the interglacial (Figure S2; Table S4). On the other hand, the temperate *Ceratoscopelus maderensis* shows a general increase in size from MIS 20 to MIS 18, whereas *Diaphus* sizes decrease.

**Figure 3.**
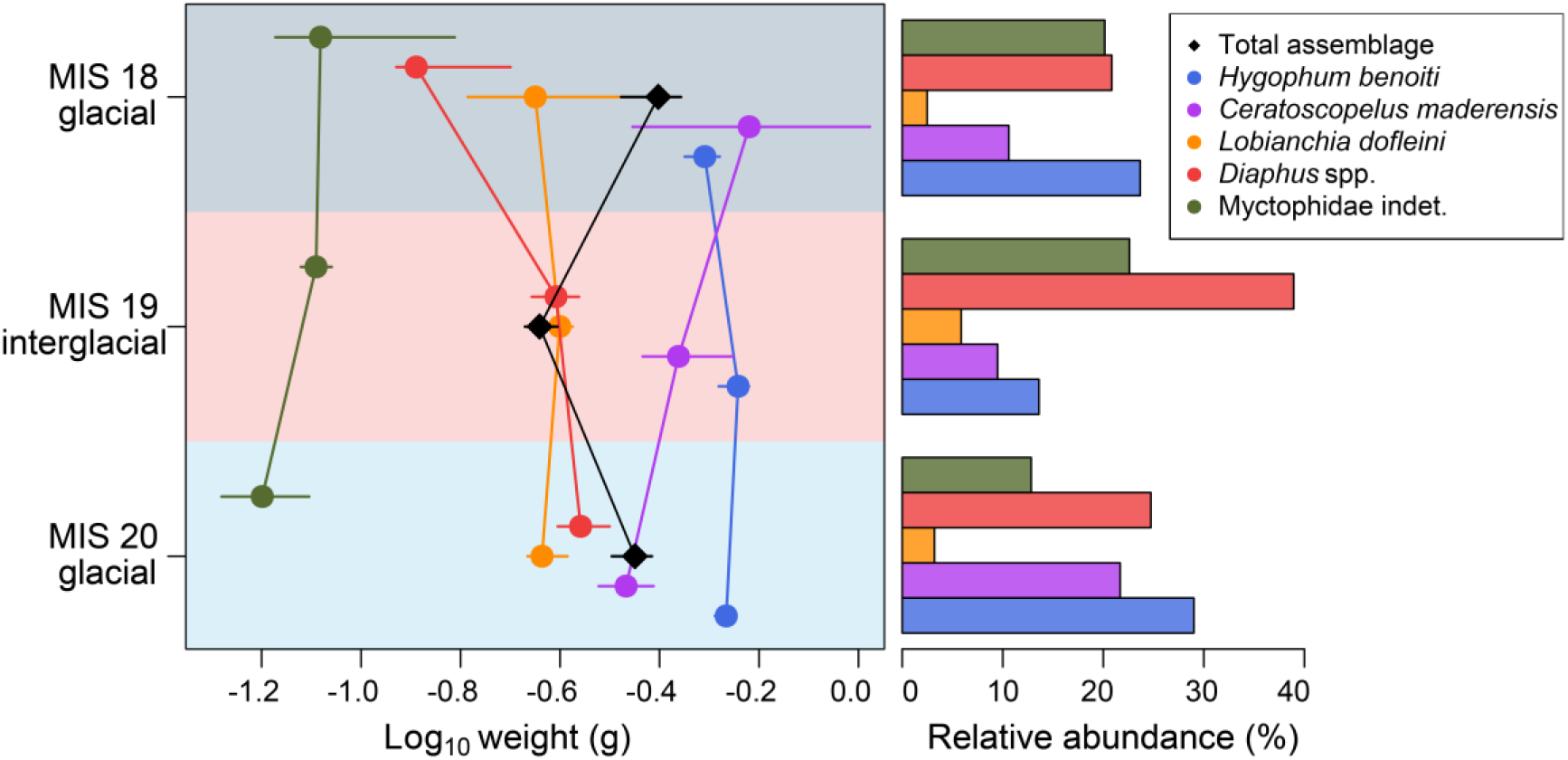
Median fish weight (with 95% bootstrapped confidence intervals) of the entire assemblage and the two most abundant cold- and warm-water species in the three studied intervals (all lanternfishes) and the corresponding abundances. Overall fish size decreases during the MIS 19 interglacial, but the individual species show different patterns.

The assemblages are clearly dominated by lanternfishes (Myctophidae constitute 89%, 96%, and 78% of the assemblages in MIS 20, MIS 19, and MIS 18, respectively; Table S4), whose median weight decreases by ~35% from 0.34 g in MIS 20 glacial to 0.22 g in MIS 19 interglacial and increase again to 0.35 g in MIS18 glacial (Kruskal-Wallis test *χ*^2^ = 51.905, df = 2, *p* < 0.001; pairwise Wilcoxon test: *p* < 0.001 for both MIS 20–19 and MIS 19–18; Figure S4). In contrast, cods (Gadidae) may show the opposite trend, with their median weight showing higher median value (2.24 g) during the interglacial, but lower in the glacials (Kruskal-Wallis test *χ*^2^ = 6.626, df = 2, *p* < 0.04; pairwise Wilcoxon test: *p* < 0.33 for MIS 20–19, *p* < 0.33 for MIS 20–18 and *p* < 0.04 for MIS 19–18). However, we cannot reach a robust conclusion about this family given the small sample sizes (9, 17 and 8 otoliths in MIS 20, 19 and 18, respectively). Other families identified in the dataset had too few specimens to identify any significant trends (Figure S4).

The median weight among cold-water species does not significantly differ between the three stages (Kuskal-Wallis test *χ*^2^ = 4.065, df = 2, *p* = 0.13; Figure S5), while the weight of warm-water species decreases from MIS 20 to MIS 19 (Kruskal-Wallis test *χ*^2^ = 16.843, df = 2, *p* < 0.001; pairwise Wilcoxon test *p* < 0.001 for MIS 20–19, *p* = 0.05 for MIS 20–18 and *p* = 0.42 for MIS 19–18). The relative abundance of cold-water (temperate and a few subpolar species) clearly drops during the MIS 19 interglacial, whereas warm-water species have a higher contribution to the MIS 19 assemblage.

Change in assemblage composition through the studied glacial–interglacial–glacial transitions is mostly driven by cold-water mesopelagic fishes (Figures 3 and S6). *Ceratoscopelus maderensis* and *Hygophum benoiti* abundances clearly drop during the MIS 20–19 deglaciation, the latter increasing again significantly in MIS 18 along with *Vinciguerria poweriae*, despite the shallowing of the study area.

The observed shifts in size and assemblage composition across the glacial–interglacial–glacial transitions cannot be explained by a variation in the otolith preservation, because the average taphonomic score of the otoliths from the three studied intervals is not significantly different (details in the supplementary information; Figures S7 & S8; Table S5).

## Discussion

The effect of past climate changes on mesopelagic fishes manifests through shifts in assemblage composition (11,12) and fish size (64). Our results show an overall drop in mesopelagic fish median length and weight from the MIS 20 glacial to the MIS 19 interglacial in the eastern Mediterranean (Figure 2), which is driven by lanternfishes that dominate the assemblages (Figures 3, S2 and S5). This decrease follows the expected effects of increasing temperature on body size of aquatic ectotherms (28,29,65), which have been observed for euphotic zone fishes today (5). The MIS 20–19 deglaciation took place within ~3,000 yrs (40,41), much longer than the scale of decades of the modern climate change. Present-day anthropogenic climate change is unprecedented, and therefore its effects are not directly comparable to those of past climate change. Nevertheless, the past can guide our understanding of the natural variability of marine ecosystems and the response of organisms to extreme environmental change. A negative relationship between climate warming and mesopelagic fish size has been observed in other areas today (66), although some studies have also shown increase in size or no relationship at all (8), and small individuals have generally been overlooked in studies of climate change impact on fish growth (67). Our results suggest that at long-term timescales, natural climate warming may lead to an overall decrease in median fish size at the community level, at least in mid-latitude regions.

Despite this overall shift, the individual myctophid genera and species in the Middle Pleistocene assemblages of the eastern Mediterranean do not follow the within-species size reduction observed in epipelagic fishes (5,31), but exhibit different and often opposing trends (Figure 3), which indicates that patterns at the assemblage level are not primarily driven by common trends in average body size of individual species. The known shifts in the fish community composition (changes in the relative abundances of species) (11) between glacials and interglacials are observed again in our study. Cold-water (temperate and subpolar) mesopelagic species show a higher relative abundance in the glacial than in the interglacial assemblages. In the Early–Middle Pleistocene Transition, North Atlantic subsurface waters experienced warming phases during glacials after MIS 24 due to pulses of increased outflow of Mediterranean waters (68), which would enhance water mixing and facilitate the functional connectivity between the mesopelagic fish populations of the two regions. This pattern may explain the establishment of subpolar species in the Mediterranean during Middle and Late Pleistocene glacials (11,12,69, 70) and the higher relative abundance of cold-water species in MIS 20 and MIS 18 observed in this study.

Moreover, the mesopelagic species may have shifted their ranges to greater, cooler depths, following their preferred temperatures, as a response to climate warming, while maintaining their body size. A case in point is *Ceratoscopelus maderensis*, whose high resilience may be explained by its large depth range, particularly in the modern Mediterranean Sea (up to 2500 m in the Ionian Sea (71)). In contrast to euphotic zone fishes, mesopelagic fishes commonly change their depth distribution. They move hundreds of meters within the water column to adapt to changing water conditions (13), and show different bathymetric distributions in different seas (72–74). This would explain size changes in individual species that may contradict expectations based on the climatic conditions (Figure 3). However, the part of the Lindos Bay Formation sedimentary sequence that was sampled here is regressive, meaning that the paleodepth decreased upwards in the section, from MIS 20 to MIS 18 (11,57). Therefore, we cannot expect shifts to greater (and colder) depths from MIS 20 to MIS 19 as an adaptation to warming climate, which would explain why the size of *Ceratoscopelus maderensis* indeed decreases in the interglacial.

In addition, temperature effects on metabolism are size-specific (74, 75). Accordingly, it is possible for fish to maintain (or even increase) their size during warming, when the increased growth (over)compensates for higher mortality in the warmer ecosystem (76–78). Size-specific impacts of temperature have been recorded in short- and long-term records, at least at the decadal scale (79). This size-specific body size responses to warming climate could be another mechanism to explain the observed unchanged median body sizes of *L. dofleini* and *H. benoiti*, and the increased size of *C. maderensis* during the Pleistocene interglacial in the eastern Mediterranean, although further studies would be required to support this.

Finally, both direct and indirect effects of elevated temperature on fishes should be taken into account to explain the divergent patterns shown by the different mesopelagic fishes. The higher temperature expedites fish growth by increasing metabolism, but at the same time may limit food availability through bottom-up effects, resulting in a decrease in growth (80). Although palaeoproductivity data for the eastern Mediterranean during MIS20–18 are not available, data from the Western Mediterranean (81) indicate increase, rather than decrease, of primary productivity during the interglacial MIS 19, thus rejecting a negative feedback of warming on fish growth at that time.

## Conclusion

Although the eastern Mediterranean mesopelagic fish assemblages seem to follow the predicted size reduction with increasing temperature during the Middle Pleistocene MIS 19 interglacial, our results demonstrate that this was due to changes in species relative abundances coupled with size-dependent temperature effects rather than on individual species body size. Within-species, synchronous reductions in size across multiple mesopelagic species were not observed in the studied assemblages, suggesting that an increase in relative abundance of small mesopelagic fishes took place in the eastern Mediterranean during MIS 19 interglacial. Even though our study refers to a climate warming taking place over a few thousand years in the Pleistocene, rather than within decades as is predicted for the modern climate change, our results are consistent with the few available observations on mesopelagic fishes today. Therefore, accurate predictions of mesopelagic fish responses under future climate change scenarios may require an ecosystem-based and multispecies rather than a single-species approach. Considering the important function of mesopelagic fishes in energy and carbon transfer through marine ecosystems, the observed community-level decrease in size of mesopelagic fishes may suggest downgrading of the marine food-web structure and a reduction in carbon sequestration during the interglacial.

## Acknowledgements

This work was funded by the Austrian Science Fund (FWF), project M2894-N “Deep-time climate change impact on marine food webs” (P.I.: KA). It was also supported by the National programs Tellus-INTERRVIE and Tellus-SYSTER of CNRS-INSU (FQ). MC acknowledges the ‘Severo Ochoa Centre of Excellence’ accreditation (CEX2019-000928-S) to the Institute of Marine Science (ICM-CSIC) and partial funding from the European Union’s Horizon 2020 research and innovation program under grant agreement No. 869300 (FutureMARES project). The authors would like to thank Norbert Frotzler for drawing the fish head used in our supplementary Figure S1. This research contributes to the objectives of Q-MARE (a PAGES working group). We would like to thank the three anonymous reviewers for the very helpful comments on our manuscript.

## Supplementary Information

https://doi.org/10.5061/dryad.gxd2547pn

## Authors Contribution

Conceptualization: KA; Formal analysis: KA, TS, RN, JF; Funding acquisition: KA & MZ; Investigation: KA, FQ, RN, TS, EK; Methodology: KA, TS, RN; Writing – original draft: KA & TS; Writing – review & editing: FQ, RN, EK, MC, MZ.

## References

1. Lotze HK, Tittensor DP, Bryndum-Buchholz A, Eddy TD, Cheung WWL, Galbraith ED, et al. Global ensemble projections reveal trophic amplification of ocean biomass declines with climate change. Proc Natl Acad Sci. 2019;201900194.

2. Pauly D. On the interrelationships between natural mortality, growth parameters, and mean environmental temperature in 175 fish stocks. ICES J Mar Sci. 1980;39(2):175–92.

3. Poloczanska ES, Burrows MT, Brown CJ, García Molinos J, Halpern BS, Hoegh-Guldberg O, et al. Responses of Marine Organisms to Climate Change across Oceans. Front Mar Sci. 2016;3:62.

4. Boyce DG, Lotze HK, Tittensor DP, Carozza DA, Worm B. Future ocean biomass losses may widen socioeconomic equity gaps. Nat Commun. 2020;11(1):2235.

5. Cheung WWL, Sarmiento JL, Dunne J, Frölicher TL, Lam VWY, Palomares MLD, et al. Shrinking of fishes exacerbates impacts of global ocean changes on marine ecosystems. Nat Clim Change. 2013;3(3):254.

6. Brito-Morales I, Schoeman DS, Molinos JG, Burrows MT, Klein CJ, Arafeh-Dalmau N, et al. Climate velocity reveals increasing exposure of deep-ocean biodiversity to future warming. Nat Clim Change. 2020;10(6):576–81.

7. Proud R, Cox MJ, Brierley AS. Biogeography of the Global Ocean’s Mesopelagic Zone. Curr Biol. 2017;27(1):113–9.

8. Thresher RE, Koslow JA, Morison AK, Smith DC. Depth-mediated reversal of the effects of climate change on long-term growth rates of exploited marine fish. Proc Natl Acad Sci. 2007;104(18):7461–5.

9. Hidalgo M, Browman HI. Developing the knowledge base needed to sustainably manage mesopelagic resources. ICES J Mar Sci. 2019;76(3):609–15.

10. Clavel-Henry M., Piroddi C., Quattrocchi F., Macias D., Christensen V. Spatial distribution and abundance of mesopelagic fish biomass in the Mediterranean Sea. Front Mar Sci. 2020;7:573986.

11. Agiadi K, Girone A, Koskeridou E, Moissette P, Cornée JJ, Quillévéré F. Pleistocene marine fish invasions and paleoenvironmental reconstructions in the eastern Mediterranean. Quat Sci Rev. 2018;196:80–99.

12. Agiadi K, Triantaphyllou M, Girone A, Karakitsios V. The early Quaternary palaeobiogeography of the eastern Ionian deep-sea Teleost fauna: A novel palaeocirculation approach. Palaeogeogr Palaeoclimatol Palaeoecol. 2011;306(3–4):228–42.

13. Catul V, Gauns M, Karuppasamy PK. A review on mesopelagic fishes belonging to family Myctophidae. Rev Fish Biol Fish. 2011;21(3):339–54.

14. Jones WA, Checkley DM. Mesopelagic fishes dominate otolith record of past two millennia in the Santa Barbara Basin. Nat Commun. 2019;10:4564.

15. Schwarzhans W, Carnevale G. The rise to dominance of lanternfishes (Teleostei: Myctophidae) in the oceanic ecosystems: a paleontological perspective. Paleobiology. 2021;47(3):446–63.

16. Saba GK, Burd AB, Dunne JP, Hernández-León S, Martin AH, Rose KA, et al. Toward a better understanding of fish-based contribution to ocean carbon flux. Limnol Oceanogr. 2021;11709

17. Langbehn TJ, Aksnes DL, Kaartvedt S, Fiksen Ø, Jørgensen C. Light comfort zone in a mesopelagic fish emerges from adaptive behaviour along a latitudinal gradient. Mar Ecol Prog Ser. 2019;623:161–74.

18. Badcock J, Merrett NR. Midwater fishes in the eastern North Atlantic—I. Vertical distribution and associated biology in 30°N, 23°W, with developmental notes on certain myctophids. Prog Oceanogr. 1976;7(1):3–58.

19. Olivar MP, Bernal A, Moli B, Pena M, Balbin R, Castellon A, Miquel J, Massuti E. Vertical distribution, diversity and assemblages of mesopelagic fishes in the western Mediterranean. Deep Sea Res I. 2012;62:53–69.

20. Murphy E j, Watkins J l, Trathan P n, Reid K, Meredith M p, Thorpe S e, et al. Spatial and temporal operation of the Scotia Sea ecosystem: a review of large-scale links in a krill centred food web. Philos Trans R Soc B Biol Sci. 2007;362(1477):113–48.

21. Anderson TR, Martin AP, Lampitt RS, Trueman CN, Henson SA, Mayor DJ. Quantifying carbon fluxes from primary production to mesopelagic fish using a simple food web model. ICES J Mar Sci. 2019;76(3):690–701.

22. Archibald KM, Siegel DA, Doney SC. Modeling the Impact of Zooplankton Diel Vertical Migration on the Carbon Export Flux of the Biological Pump. Glob Biogeochem Cycles. 2019;33(2):181–99.

23. Aumont O, Maury O, Lefort S, Bopp L. Evaluating the Potential Impacts of the Diurnal Vertical Migration by Marine Organisms on Marine Biogeochemistry. Glob Biogeochem Cycles. 2018;32(11):1622–43.

24. Pauly D, Piroddi C, Hood L, Bailly N, Chu E, Lam V, et al. The Biology of Mesopelagic Fishes and Their Catches (1950–2018) by Commercial and Experimental Fisheries. J Mar Sci Eng. 2021;9(10):1057.

25. St. John MA, Borja A, Chust G, Heath M, Grigorov I, Mariani P, et al. A Dark Hole in Our Understanding of Marine Ecosystems and Their Services: Perspectives from the Mesopelagic Community. Front Mar Sci. 2016;3:31.

26. Irigoien X, Klevjer TA, Røstad A, Martinez U, Boyra G, Acuña JL, et al. Large mesopelagic fishes biomass and trophic efficiency in the open ocean. Nat Commun. 2014;5(1):3271.

27. Hildrew AG, Raffaelli DG, Edmonds-Brown R, editors. Body Size: The Structure and Function of Aquatic Ecosystems [Internet]. Cambridge: Cambridge University Press; 2007.

28. Daufresne M, Lengfellner K, Sommer U. Global warming benefits the small in aquatic ecosystems. Proc Natl Acad Sci. 2009;106(31):12788–93.

29. Ohlberger J. Climate warming and ectotherm body size – from individual physiology to community ecology. Funct Ecol. 2013;27(4):991–1001.

30. Audzijonyte A, Fulton E, Haddon M, Helidoniotis F, Hobday AJ, Kuparinen A, et al. Trends and management implications of human-influenced life-history changes in marine ectotherms. Fish Fish. 2016;17(4):1005–28.

31. Baudron AR, Needle CL, Rijnsdorp AD, Marshall CT. Warming temperatures and smaller body sizes: synchronous changes in growth of North Sea fishes. Glob Change Biol. 2014;20(4):1023–31.

32. Dietl GP, Kidwell SM, Brenner M, Burney DA, Flessa KW, Jackson ST, et al. Conservation Paleobiology: Leveraging Knowledge of the Past to Inform Conservation and Restoration. Annu Rev Earth Planet Sci. 2015;43(1):79–103.

33. Lisiecki LE, Raymo ME. Plio–Pleistocene climate evolution: trends and transitions in glacial cycle dynamics. Quat Sci Rev. 2005;26(1):56–69.

34. Lisiecki LE, Raymo ME. A Pliocene-Pleistocene stack of 57 globally distributed benthic δ18O records. Paleoceanography. 2005;20(1).

35. McClymont EL, Sosdian SM, Rosell-Melé A, Rosenthal Y. Pleistocene sea-surface temperature evolution: Early cooling, delayed glacial intensification, and implications for the mid-Pleistocene climate transition. Earth-Sci Rev. 2013;123:173–93.

36. González-Donoso JM, Serrano F, Linares D. Sea surface temperature during the Quaternary at ODP Sites 976 and 975 (western Mediterranean). Palaeogeogr Palaeoclimatol Palaeoecol. 2000;162(1):17–44.

37. Ruddiman WF, Raymo ME, Martinson DG, Clement BM, Backman J. Pleistocene evolution: Northern hemisphere ice sheets and North Atlantic Ocean. Paleoceanography. 1989;4(4):353–412.

38. Marino M, Girone A, Gallicchio S, Herbert T, Addante M, Bazzicalupo P, et al. Climate variability during MIS 20–18 as recorded by alkenone-SST and calcareous plankton in the Ionian Basin (central Mediterranean). Palaeogeogr Palaeoclimatol Palaeoecol. 2020;560:110027.

39. Quivelli O, Marino M, Rodrigues T, Girone A, Maiorano P, Bertini A, et al. Multiproxy record of suborbital-scale climate changes in the Algero-Balearic Basin during late MIS 20 - Termination IX. Quat Sci Rev. 2021;260:106916.

40. Nomade S, Bassinot F, Marino M, Simon O, Dewilde F, Maiorano P, Isguder G, Blamart D, Girone A, Scao V, Pereira A, Toti F, Bartini A, Combourieu-Nebout N, Peral M, Bourles DL, Petrosino P, Gallicchio S, Ciaranfi N. High-resolution foraminifer stable isotope record of MIS19 at Montalbano Jonico, southern Italy: A window into Mediterranean climatic variability during a low-eccentricity interglacial. Quat Sci Rev. 2019;205:106–125.

41. Simon Q, Bourles DL, Bassinot F, Nomade S, Marino M, Ciaranfi N, Girone A, Maiorano P, Thouveny N, Choy S, Dewilde F, Scao V, Isguder G, Blamart D. Authigenic 10Be/9Be ratio signature of the Matuyama–Brunhes boundary in the Montalbano Jonico marine suggession. Earth Planet Sci Letters. 2017;460:255–267.

42. Cheung WWL, Reygondeau G, Frölicher TL. Large benefits to marine fisheries of meeting the 1.5°C global warming target. Science. 2016;354(6319):1591–4.

43. IPCC. Climate Change 2021: The Physical Science Basis. Contribution of Working Group I to the Sixth Assessment Report of the Intergovernmental Panel on Climate Change. [Masson-Delmotte, V., P. Zhai, A. Pirani, S. L. Connors, C. Pean, S. Berger, N. Caud, Y. Chen, L. Goldfarb, M. I. Gomis, M. Huang, K. Leitzell, E. Lonnoy, J. B. R. Matthews, T. K. Maycock, T. Waterfield, O. Yelekci, R. Yu and B. Zhoo (eds.)]. Cambridge University Press; 2021.

44. IPCC. Special Report on the Ocean and Cryosphere in a changing climate. 2019 p. 765. (Working Group II. Technical support unit). Available from: https://www.ipcc.ch/site/assets/uploads/sites/3/2019/12/SROCC_FullReport_FINAL.pdf

45. Nolf D. Otolithi Piscium. Schlutze H.P., editor. Stuttgart: G. Fischer Verlag; 1985. 145 p. (Handbook of paleoichthyology; vol. 10).

46. Agiadi K, Albano PG. Holocene fish assemblages provide baseline data for the rapidly changing eastern Mediterranean: The Holocene. 2020;30(10):1438–50.

47. Lin CH, Gracia BD, Pierotti MER, Andrews AH, Griswold K, O’Dea A. Reconstructing reef fish communities using fish otoliths in coral reef sediments. PLOS ONE. 2019;14(6):e0218413.

48. Edelist D. New length–weight relationships and Lmax values for fishes from the Southeastern Mediterranean Sea. J Appl Ichthyol. 2014;30(3):521–6.

49. Mosegaard H, Svedäng H, Taberman K. Uncoupling of Somatic and Otolith Growth Rates in Arctic Char (Salvelinus alpinus) as an Effect of Differences in Temperature Response. Can J Fish Aquat Sci. 1988;45(9):1514–24.

50. Cornée JJ, Quillévéré F, Moissette P, Fietzke J, López-Otálvaro GE, Melinte-Dobrinescu M, et al. Tectonic motion in oblique subduction forearcs: insights from the revisited Middle and Upper Pleistocene deposits of Rhodes, Greece. J Geol Soc. 2019;176(1):78–96.

51. Quillévéré F, Cornée JJ, Moissette P, López-Otálvaro GE, van Baak C, Münch P, et al. Chronostratigraphy of uplifted Quaternary hemipelagic deposits from the Dodecanese island of Rhodes (Greece). Quat Res. 2016;86(1):79–94.

52. Van Hinsbergen DJJ, Krijgsman W, Langereis CG, Cornée JJ, Duermeijer CE, Van Vugt N. Discrete plio-Pleistocene phases of tilting and counterclockwise rotation in the southeastern Aegean arc (Rhodos, Greece): Early Pliocene formation of the south Aegean left-lateral strike-slip system. J Geol Soc. 2007;164(6):1133–44.

53. Quillévéré F, Nouailhat N, Joannin S, Cornée JJ, Moissette P, Lécuyer C, et al. An onshore bathyal record of tectonics and climate cycles at the onset of the Early-Middle Pleistocene Transition in the eastern Mediterranean. Quat Sci Rev. 2019;209:23–39.

54. Moissette P, Cornée JJ, Quillévéré F, Zibrowius H, Koskeridou E, López-otálvaro G emperatriz. Pleistocene (Calabrian) deep-water corals and associated biodiversity in the eastern Mediterranean (Karpathos Island, Greece). J Quat Sci. 2017;32(7):923–33.

55. van Hinsbergen DJJ, Meulenkamp JE. Neogene supradetachment basin development on Crete (Greece) during exhumation of the South Aegean core complex. Basin Res. 2006;18(1):103–24.

56. Balmer EM, Robertson AHF, Raffi I, Kroon D. Pliocene–Pleistocene sedimentary development of the syntectonic Polis graben, NW Cyprus: evidence from facies analysis, nannofossil biochronology and strontium isotope dating. Geol Mag. 2018;1–29.

57. Titschack J, Joseph N, Fietzke J, Freiwald A, Bromley RG. Record of a tectonically-controlled regression captured by changes in carbonate skeletal associations on a structured island shelf (mid-Pleistocene, Rhodes, Greece). Sediment Geol. 2013;283(Supplement C):15–33.

58. Froese R, Thorson JT, Reyes RB. A Bayesian approach for estimating length-weight relationships in fishes. J Appl Ichthyol. 2014;30(1):78–85.

59. Giménez J, Manjabacas A, Tuset VM, Lombarte A. Relationships between otolith and fish size from Mediterranean and north-eastern Atlantic species to be used in predator–prey studies. J Fish Biol. 2016;89(4):2195–202.

60. Kaschner KK, Kesner-Reyes K, Garilao C, Rius-Barile J, Rees T, Froese R. AquaMaps: predicted range maps for aquatic species. www.aquamaps.org, version 08/2016; 2016.

61. Froese R, Pauly D. FishBase. 2022. Available from: www.fishbase.org

62. Agiadi K, Azzarone M, Hua Q, Kaufman D, Thivaiou D, Albano PG. The taphonomic clock in fish otoliths. Paleobiology. 2021;https://doi.org/10.1017/pab.2021.30

63. Agiadi K, Nawrot R, Albano PG, Koskeridou E, Zuschin M. Potential and limitations of applying the mean temperature approach to fossil otolith assemblages. Environ Biol Fishes. 2022;https://doi.org/10.1007/s10641-022-01252-6

64. Salvatecci R, Schneider RR, Galbraith E, Field DB, Blanz T, Bauersachs T, et al. Smaller fish species in a warm and oxygen-poor Humboldt Current system. Science. 2022;375:101–4.

65. Bergmann C. Über die Verhältnisse der Wärmeökonomie der Thiere zu ihrer Grösse. Gött Stud. 1847;1:595–708.

66. Saunders RA, Tarling GA. Southern Ocean Mesopelagic Fish Comply with Bergmann’s Rule. Am Nat. 2018 Mar 1;191(3):343–51.

67. Huang M, Ding L, Wang J, Ding C, Tao J. The impacts of climate change on fish growth: A summary of conducted studies and current knowledge. Ecol Indic. 2021;121:106976.

68. Catunda MCA, Bahr A, Kaboth-Bahr S, Zhang X, Foukal NP, Friedrich O. Subsurface Heat Channel Drove Sea Surface Warming in the High-Latitude North Atlantic During the Mid-Pleistocene Transition. Geophys Res Lett. 2021;48(11):e2020GL091899.

69. Girone A, Nolf D, Cappetta H. Pleistocene fish otoliths from the Mediterranean Basin: a synthesis. Geobios. 2006;39(5):651–71.

69. Lin CH, Taviani M, Angeletti L, Girone A, Nolf D. Fish otoliths in superficial sediments of the Mediterranean Sea. Palaeogeogr Palaeoclimatol Palaeoecol. 2017;471:134–43.

70. Mytilineou C, Politou CY, Papaconstantinou C, Kavadas S, D’Onghia G, Sion L. Deep-water fish fauna in the Eastern Ionian Sea. Belg J Zool. 2005;135(2):229–33.

71. Olivar MP, Bernal A, Molí B, Peña M, Balbín R, Castellón A, et al. Vertical distribution, diversity and assemblages of mesopelagic fishes in the western Mediterranean. Deep Sea Res Part Oceanogr Res Pap. 2012;62:53–69.

72. Badcock J, Merrett NR. Midwater fishes in the eastern North Atlantic—I. Vertical distribution and associated biology in 30°N, 23°W, with developmental notes on certain myctophids. Prog Oceanogr. 1976;7(1):3–58.

73. Kinzer J, Schulz K. Vertical distribution and feeding patterns of midwater fish in the central equatorial Atlantic. Mar Biol. 1985;85(3):313–22.

74. Killen SS, Atkinson D., Glazier DS. The intraspecific scaling of metabolic rate with body mass in fishes depends on lifestyle and temperature. Ecol Lett. 2010;13:184–93.

75. Ohlberger J, Mehner T, Staaks G, Hölker F. Intraspecific temperature dependence of the scaling of metabolic rate with body mass in fishes and its ecological implications. Oikos. 2012;121:245–51.

76. Ohlberger J, Edeline E, Vollestad LA, Stenseth NC, Claessen D. Temperature-driven regime shifts in the dynamics of size-structured populations. Am Nat. 2011;177:211–23.

77. Ohlberger J, Langangen Ø, Edeline E, Claessen D, Winfield IJ, Stenseth NC, Vøllestad LA. Stage-specific biomass overcompensation by juveniles in response to increased adult mortality in a wild fish population. Ecology. 2011;92:2175–82.

78. Lindmark M, Huss M, Ohlberger J, Gårdmark A. Temperature-dependent body size effects determine population responses to climate warming. Ecol Lettr. 2018;21(2):181–9.

79. Huss M, Lindmark M, Jacobson P, Renee M v D, Gårdmark A. Experimental evidence of gradual size-dependent shifts in body size and growth of fish in response to warming. Global Change Biol. 2019;25(7):2285–95.

80. Lindmark M, Audzijonyte A, Blachard JL, Gårdmark A. Temperature impacts on fish physiology and resource abundance lead to faster growth but smaller fish sizes and yields under warming. Global Change Biol. 2022;28(21):6239–53.

81. Marino M, Rodrigues T, Quivelli O, Girone A, Maiorano P, Bassinot F. Paleoproductivity proxies and alkenone precursors in the Western Mediterranean during the Early–Middle Pleistocene transition. Palaeogeogr Palaeocl Palaeoecol. 2022;601:111104.

